# Characterization of Gait Kinematics and Muscle Function in Becker Muscular Dystrophy Pigs: a pilot study

**DOI:** 10.1101/2025.10.10.681620

**Authors:** Aaron B. Morton, Jacob A. Kendra, Shadi Golpasandi, Macie Mackey, Haley Russell, Connor Hendrie, Kuanting Chen, Zoe E. Kiefer, Jennifer M. Yentes, Jason W. Ross, Joshua T. Selsby, Nicolaas E. P. Deutz, Peter P. Nghiem

## Abstract

Vertebrate animal models of Becker muscular dystrophy (BMD) have been developed. Here, we characterized the gait kinematics and muscle function of a naturally occurring BMD pig model of dystrophin insufficiency. BMD pigs tended to have alterations in hip range of motion (ROM): hip (67%, 95% CI -0.64 to 14.12 degrees). While parameters were unaltered in extensor muscles, the dystrophin levels in flexor tibiotarsal joint muscles correlated with fatigue index as well as reduced isometric force (48%, 95% CI -1.86 to -0.61 N-m), and a 33% increase in fatigue index (95% CI -36.25 to 96.71 percent); the extensor muscles had no observable reductions in muscle force, with a 48% increase in fatigue index (95% CI -232.6 to 472.6 percent). Histological analysis of muscle biopsies supported a BMD phenotype in the flexor muscles of BMD pigs, with a 75% (95% CI -55.14 to -15.66 percent) decrease in large and a 43% (95% CI 17.74 to 57.38 percent) increase in small muscle fiber cross-sectional area. Dystrophin protein abundance was 28% less in flexor muscles from BMD pigs (95% CI -49.63 to 11.41 arbitrary units). Together, our model may serve as a clinically relevant model of BMD to assess safety and efficacy of therapeutics.

## Introduction

Becker muscular dystrophy (BMD) describes a spectrum of rare X-linked genetic disease caused by mutations in the *DMD* gene, resulting in the production of a truncated and/or insufficient accumulation of functional dystrophin protein for 1 in 18,000 male births (Gorgoglione et al., 2025; Magot et al., 2023; Malhotra et al., 1988). As the milder allelic variant of Duchenne muscular dystrophy (DMD), BMD typically presents later in childhood/adolescence, progressing slower with reduced severity, though symptoms are widely heterogenous among affected populations (Le Rumeur, Winder, & Hubert, 2010; Mercuri, Bönnemann, & Muntoni, 2019). Common phenotypes of BMD patients include progressive muscle weakness and degeneration of the pelvic and shoulder girdle, gait abnormalities, and an increased risk of cardiomyopathy (Flanigan, 2014; Salari et al., 2022). Despite the clinical significance, BMD remains poorly understood compared to DMD, with limited therapeutic development and fewer dedicated clinical studies (Straub & Guglieri, 2023). Given the prominence of gene and exon skipping therapies aimed to lessen the severity of DMD to a Becker-like phenotype (Duan, 2018; Straub & Guglieri, 2023), there is a need to develop translatable preclinical models of BMD.

Several DMD animal models have been developed for therapeutic testing, including multiple murine models, rodent, dogs, and pigs, but only recently was a murine model characterized for BMD (Blasi et al., 2025; Wilson et al., 2017). A porcine model of BMD would possess several advantages over rodent and dog models as their anatomy, physiology, and genome more similarly mimic those of humans (Hollinger et al., 2014; Stirm, Klymiuk, Nagashima, Kupatt, & Wolf, 2024).

Our group has characterized the *DMD* gene in-frame mutation of a naturally occurring BMD porcine model that appears to be caused by a nucleotide change in Exon 41, which disrupts the amino acid sequence resulting in an insufficient accumulation of the dystrophin protein (Hollinger et al., 2014; Nonneman, Brown-Brandl, Jones, Wiedmann, & Rohrer, 2012; Selsby, Ross, Nonneman, & Hollinger, 2015). At 12 months of age, these BMD pigs had reduced dystrophin expression in the diaphragm (70% reduction), the longissimus (90% reduction), and the psoas (45% reduction) compared to healthy littermates, as well as varied muscle histopathology (Hollinger et al., 2014; Selsby et al., 2015). Furthermore, preliminary investigations in step length and step time demonstrated gait abnormalities indicative of disease-related compensation and elevated creatine kinase (CK) levels (Selsby et al., 2015). We also observed cardiomyopathy and electrocardiogram abnormalities (Selsby et al., 2015). A heterogenous phenotype is observed across investigated skeletal muscles, likely arising from differences in the quadruped gait cycle. The initial, limited characterization of this model suggests some similar disease manifestations as human BMD patients, yet further investigations are required to characterize this model as an effective large animal, preclinical model for therapeutic safety and efficacy testing prior to human clinical trials.

The purpose of this study was to further investigate characteristics of a naturally occurring BMD porcine model of dystrophin insufficiency. We assessed hindlimb gait kinematics, hindlimb muscle function, histopathology, and blood chemistry in Healthy and BMD (Affected) NIH minipigs, 8-10 months of age. Herein, we report further abnormalities in the BMD pig hindlimb muscles, providing additional insight into the pathology of a large animal, preclinical model of BMD.

## Materials and Methods

### Ethical Statement

All protocols in this study were approved by Animal Care and Use Committees at Texas A&M University (AUP 2023-0025). Animal care was in accordance with the National Research Council’s *Guide for the Care and Use of Laboratory Animals (Eighth Edition, 2011)*. Animals were coded prior to experimentation and data analysis was conducted by blind investigators.

### Animals

Male and Female NIH minipigs (Healthy, n=2 and Affected, n=5) were bred at Iowa State University and transported to Texas A&M University and maintained in approved kennels. Boars were single housed to prevent fighting but were able to visually see other animals in the bay. Animal numbers were limited by age and availability.

### Surgical Muscle Biopsy

Animals were sedated with butorphanol, masked, and maintained with inhalant isoflurane maintained at 2% unless an increase was required up to 4% and 100% oxygen. Following blood collection, the surgical site was aseptically prepped. A 3-4 cm incision was made in the skin and a 1×1×1cm open muscle biopsy was taken from the vastus lateralis, biceps femoris, cranial tibial (flexor), and lateral head of the gastrocnemius (extensor). The incision was closed with 2-0 Vicryl, the animal recovered from anesthesia and returned to its home cage with post operative monitoring and analgesic as needed.

### Isometric muscle testing

Methods were adapted from our internationally established canine protocols and modified based on published torque measures in pigs ( Blasi et al., 2025; Corona, Rivera, Dalske, Wenke, & Greising, 2020). Animals were placed under general anesthesia as indicated above. An Aurora Scientific, custom large animal testing machine was used for analysis (model # 701C for high phase stimulator; model # 892A-HD for the custom muscle testing rig, Aurora, Ontario, CA,). A pelvic limb was secured to a leg brace and the hooves/tarsus secured to a foot pedal with Vet Wrap (Marietta, GA). The foot pedal was connected to a transducer (Model # 492A-AMP). Percutaneous needle electrodes were inserted around the fibular nerve to stimulate (100 Hz, 0.1ms pulse, over 800 millisecond) the cranial tibial compartment to flex (pull the foot pedal) the tibiotarsal joint (90 degrees). The same stimulation was applied to the tibial nerve to stimulate the gastrocnemius and digital flexors to extend (push the foot pedal) the tibiotarsal joint (90 degrees). Twitch and tetanic contractions were performed in each muscle group to evaluate torque. All assessments were performed in flexion before moving to extension or vice versa. Due to limitations in the hardware and software, eccentric contraction decrements could not be made. A fatigue protocol was developed that delivered one tetanic contraction (100 Hz) per second for 60 replicates at 800 ms train durations with 1 second in between each replicate to induce fatigue in each muscle group. Data were recorded using DMA Software for 615A model number. Area under the curve (AUC) of fatigue replicates was analyzed between groups. Fatigue index (FI) was calculated as : FI = [(initial force – final force)/initial force) x 100].

### Gait Analysis

Beginning at 4-5 months of age, pigs underwent 2-3 months of acclimation training on a treadmill at a speed of 1 m/sec prior to motion capture and video recording. A custom treadmill with enclosed sides was used for both training and testing. The right side of the treadmill featured a transparent plexiglass panel, enabling the capture of gait kinematics. Due to the equipment set-up, the treadmill had a constant incline of 3 degrees during all walking trials. A digital video camera (GoPro 11 Black, GoPro, Inc., San Mateo, CA, U.S.) was positioned perpendicular to the treadmill to record the two-dimensional kinematics of the pigs’ gait (**Figure 1**). The camera was 1.4 meters from the treadmill at a height that centered the treadmill within the frame. To minimize interlacing and lens distortion, video recording was captured at 60 frames per second with 4K resolution, using the “Linear + Horizon Lock” setting on GoPro. Two additional lights were positioned near the treadmill at a 45-degree angle to improve visibility of the pig’s hind legs (**Figure 1**). A yardstick was placed on the treadmill parallel to the belt for image calibration.

**Figure 1.**
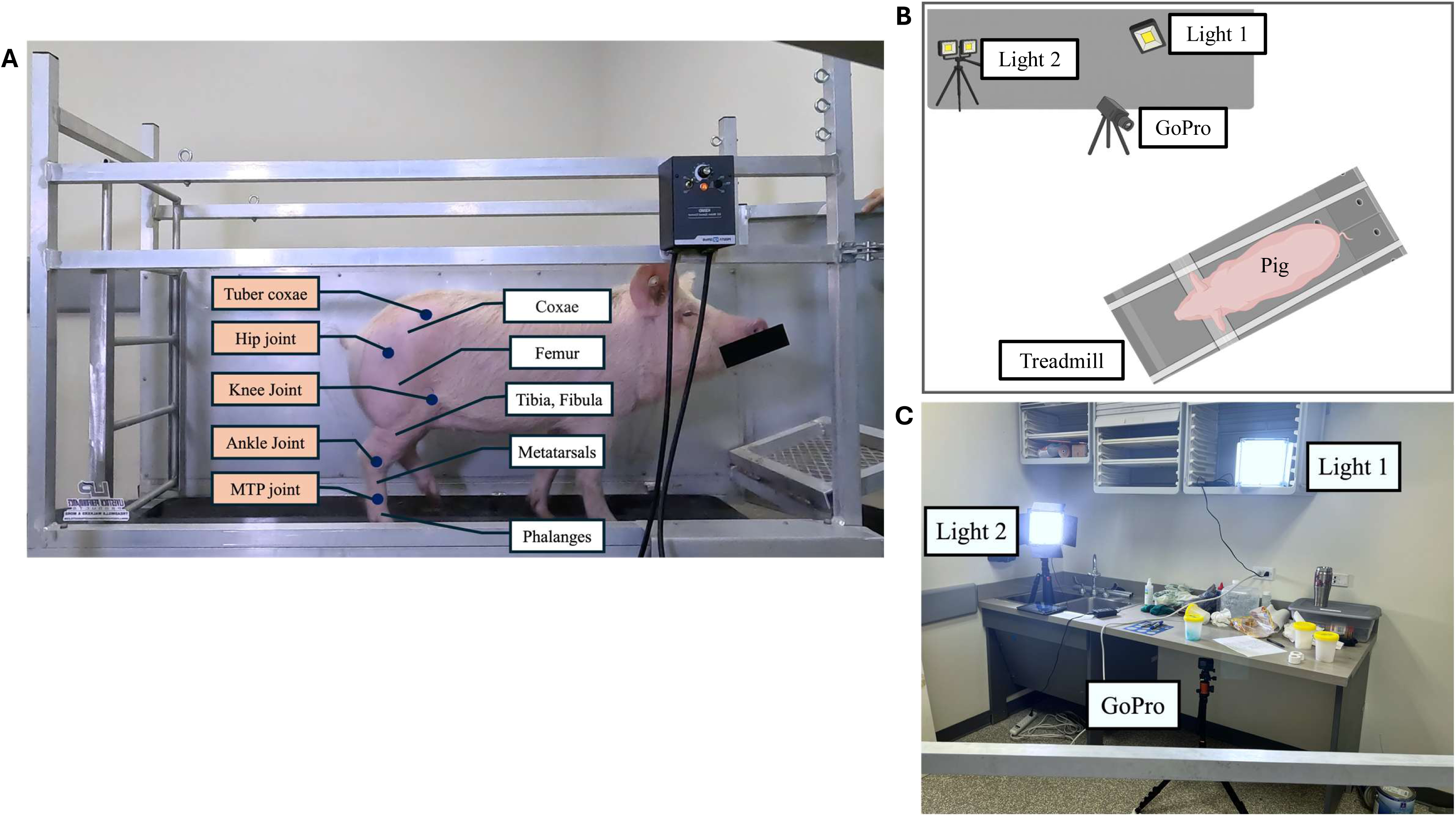
Experimental set up and marker placement for motion capture analysis. A) *Marker Placement*, Markers were placed at joints (tuber coxae, hip joint, knee joint, ankle joint, mtp joint) on the right hindlimb of walking pigs. B and C) *Room Layout*, Two lights were placed on either side of a digital camera mounted upon a tripod placed at a 90° angle to the joint markers placed upon pigs, B) Top View, C) Subject View.

Pigs’ right pelvic limb and pelvis were modeled into four distinct segments: pelvis, femur, shank, and foot using five anatomical markers placed on tuber coxae, greater trochanter, knee joint, tibiotarsal joint, and metatarsophalangeal joint. Specifically, pelvis segment was defined by markers on the tuber coxae and greater trochanter; femur was defined by markers on the greater trochanter and knee joint; shank segment was defined by markers on the knee joint and tibiotarsal joint; and the foot segment by markers on the tibiotarsal joint, and metatarsophalangeal joint. One day before data collection, a veterinarian palpated these anatomical landmarks and outlined each landmark by drawing a one-inch diameter circle using a black Sharpie. Hair around these areas was shaved to enhance marker visibility.

### Procedure

Gait data collection was conducted between 9:00 to 11:00 am. Pigs were transported individually via enclosed cart from their housing units to the treadmill located on the same hallway. The animal handler ensured that the pig was positioned at the center of the treadmill before gradually increasing the speed of the belt to 1 m/sec. Once a stable walking pattern was observed, a research technician initiated video recording. Each pig was recorded for one minute, yielding more than 60 steps for gait analysis. After recording, the pig was returned to its housing unit in an adjacent room.

### Data Analysis

Raw video recordings were exported from the GoPro camera and imported into Kinovea (v. 2023.1, Kinovea.org) for two-dimensional coordinate data extraction. Video image calibration was performed using a yardstick placed on the treadmill. A coordinate frame was established by placing the origin at the bottom left corner of the treadmill in the video. The coordinates of all five markers were extracted relative to this origin.

Marker positions were determined using a semi-automatic tracking approach. In the first video frame, a square area containing the marker was selected as a reference feature for Kinovea’s point tracking tool. This tool identified marker positions in subsequent frames by comparing pixel similarity across frames. After automatic tracking, a research technician visually inspected and manually corrected any tracking deviations. A second research technician independently reviewed the tracking results to ensure accuracy and consistency. For each recording, two-dimensional marker position time series data of the five markers were exported to Matlab (v. 2024a, MathWorks, Natick, MA) for further kinematic analysis.

Steps were classified as non-walking steps when irregular walking movements, such as kicking or jumping, were identified through comparison with the video recordings, and removed. For all remaining steps, comprehensive gait characteristics were quantified using spatiotemporal parameters and joint angle measurements. Spatiotemporal parameters included step time, stance time, swing time, swing and stance phase percentages within a step cycle, and step length.

Stride was defined as described (Boakye et al., 2020): a stride was the interval between two consecutive hoof contacts with the same hoof; the stance phase was between initial hoof contact to hoof lift-off; and the swing phase extended from hoof lift-off to the next hoof contact. Joint angles were computed based on vectors formed by two adjacent limb segments. To enable step-to-step comparisons, joint angle data were normalized for time to 100 data points per step. All joint angle plots were visually inspected to exclude non-walking steps from the final analysis. Joint angle measurements exported included maximum, minimum, and range of motion (ROM) for the hip, knee, and ankle joints.

Standard deviations from each gait analysis parameter were converted to a percentage of the healthy group and then collectively pooled into healthy and affected groups to draw conclusions around gait consistency.

### Cryosectioning, Hematoxylin and Eosin Staining, and Light Microscopic Imaging

Cranial tibial and gastrocnemius muscle samples were taken by biopsy from BMD and wild-type pigs, immediately frozen in isopentane cooled in liquid nitrogen, and stored at -80°C. Frozen skeletal muscle samples were mounted in Tissue-Plus^TM^ O.C.T. Compound (Scigen, Fischer Scientific, Hampton, NJ, USA) and sectioned (thickness, 10 μm) in a Cryostar NX50 Cryostat (Epredia, Kalamazoo, MI, USA) at -17°C onto a microscope slide, followed by H&E staining (SelecTech H&E Staining System, Leica Biosystems, Deer Park, IL, USA). Tissues were stained in Hematoxylin for 30 seconds, washed in tap water, differentiated in 0.3% acetic acid for 30 seconds, blueing agent for 1 minute and 30 seconds, then washed again in tap water. Sections were then washed with ethanol at 70%, 95%, 95% for 1 minute each, stained with Eosin for 2 minutes, and washed again with 95% and 100% ethanol. The sections were cleared in xylene and mounted using Permount^TM^ Mounting Medium (Electron Microscopy Sciences, Hatfield, PA, USA). Images were captured with a 10x (N.A., 0.50) objective using a Zeiss Axioplan upright microscope equipped with a Zeiss Axiocam HRc brightfield microscope camera (Zeiss Microscopy, White Plains NY, USA). Myofiber cross-sectional area (CSA) was analyzed from approximately 350 fibers per muscle using NIH ImageJ (RRID: SCR_003070, NIH, Bethesda, MD, USA) as described (A. B. Morton et al., 2018).

### Creatine Kinase and Biochemical Analysis

Creatine kinase and analytes (Total Protein, Albumin, Calcium, Phosphorus, Glucose, BUN, Creatinine, Bilirubin, ALKP, AST, ALT, Globulins, Albumin: Globulin ratio, Sodium, Potassium, Chloride, Sodium: Potassium ratio) were determined on a blood chemistry panel via colorimetric assays, ion selective electrodes (ISE) (Na, K, and Cl), and calculations (globulins, A:G ratio, Na:K ratio) on a Beckman Coulter DxC 700 AU analyzer through an AAVLD-accredited laboratory.

### Western Blot Analysis

Cranial tibial and gastrocnemius biopsy samples from BMD and wild-type porcine were homogenized in lysis buffer (pH 6.8) containing 75 mM Tris-HCI, 10% SDS, 10 mM EDTA, and 5% 2-mercaptoethanol as described (Fiorillo et al., 2015). Homogenates were centrifuged at 10,000 *g* for 10 minutes at 4 °C. The supernatant (soluble fraction) was aspirated, and its protein concentration assessed using the Bradford method (Sigma-Aldrich, St. Louis, MO, USA). Protein samples (50 µg) were blotted and analyzed using Image Studio Lite as described (A. Morton et al., 2025). Primary antibody used is a 1:1 mix of DYS-1 (1:1000, RRID: AB_442080) + DYS-2 (1:1000, RRID: AB_442081) monoclonal antibodies against dystrophin (Leica Biosystems, Deer Park, IL, USA). Membrane was exposed to IRDye 800CW Goat anti-Mouse IgG (1:5000, RRID: AB_621842, Cat.# 926-32210) (Li-Cor Biotechnology, Lincoln, NE, USA) and scanned with a Li-Cor Odyssey DLx Imager (Li-Cor Biotechnology, Lincoln, NE, US). Blots were normalized to total protein (A.U.) using Revert total protein stain (Cat.# 926-11011, Li-Cor Biotechnology, Lincoln, NE, USA). Results reported as percent of Healthy.

### Analysis

Summary data are reported as percent change and 95% CI from non-parametric T-Tests. Given limited numbers of healthy pig littermates, correlations were generated combining all 7 pigs using JASP (University of Amsterdam, Amsterdam, The Netherlands) and plotting individual outcome measures from each pig against Dystrophin 1 & 2 expression from both flexor and extensor muscles before analyzing respective slopes for deviation from 0. *P* ≤ 0.05 was considered statistically significant.

## Results

### General Measures

Gait was 30% less consistent in Affected pigs over Healthy (95% CI 3.68 to 54.4) **(Figure 2A)** while body weights at 9 months of age were similar between Healthy and Affected pigs **(Figure 2B),** means 50.6 kg (95% CI 49.9 to 51.3) and 50 kg (95% CI 42.2 to 57.2) respectively, (n=2-5/group).

**Figure 2.**
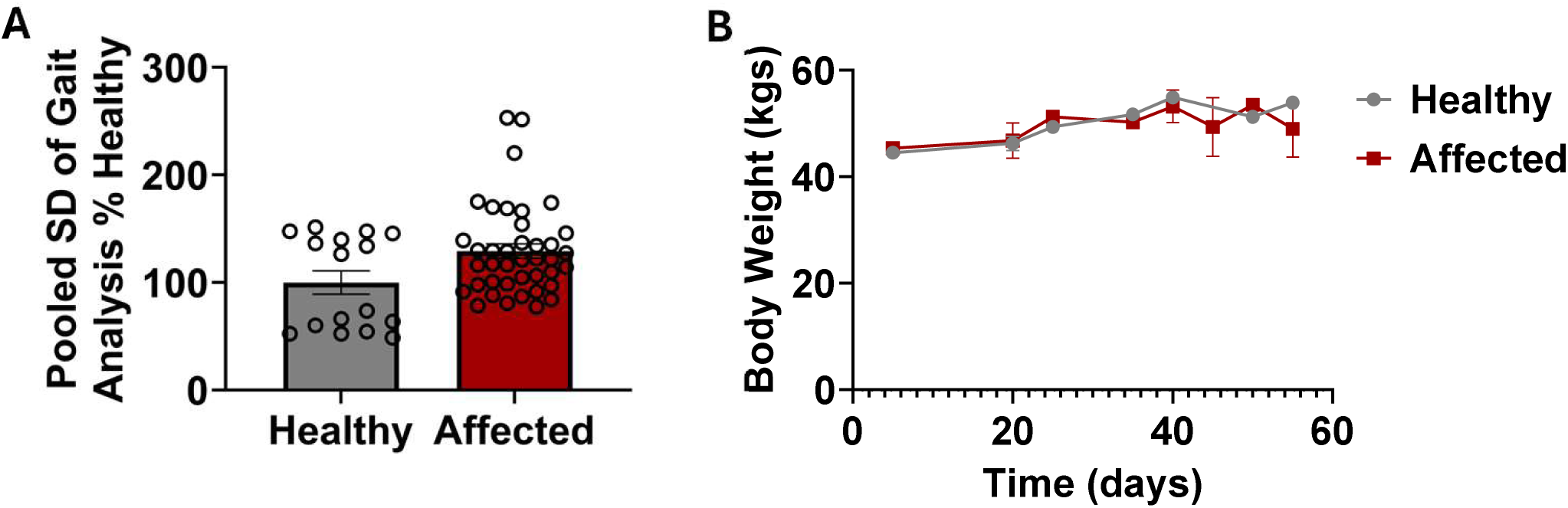
General descriptive measures. A) Pooled standard deviation (SD) reported as percent Healthy from all kinematic measures reported n=16 measures from Healthy and n=40 measures for Affected suggesting step pattern inconsistency. AB Average body weight of Healthy and Affected pigs at 9 months of age were similar. C) Body weight changes at 8-9 months of age were similar.

### Gait Analysis

Gait results demonstrated alterations to spatiotemporal and joint angle kinematics in Affected pigs compared to Healthy **(Figure 3)**. Spatiotemporal analysis of the pig gait cycles revealed swing time was decreased 12% (95% CI -0.08 to 0.01), stance time was decreased 17% (95% CI -0.16 to 0.01), and step time was decreased 15% (95% CI -0.24 to 0.01) in Affected pigs compared to Healthy **(Table 1)**. Notably, joint angle analysis demonstrated that Hip ROM and SD were increased by 40% (CI -0.64 to 14.12) and 50% (CI -0.03 to 1.67), respectively, in Affected compared to Healthy pigs **(Figure 3A)**, whereas Knee (8%, CI -17.14 to 12.13) **(Figure 3B)** and Ankle (11%, CI -15.88 to 8.46) ROM declined slightly in Affected compared to Healthy, respectively **(Figure 3C)**.

**Figure 3.**
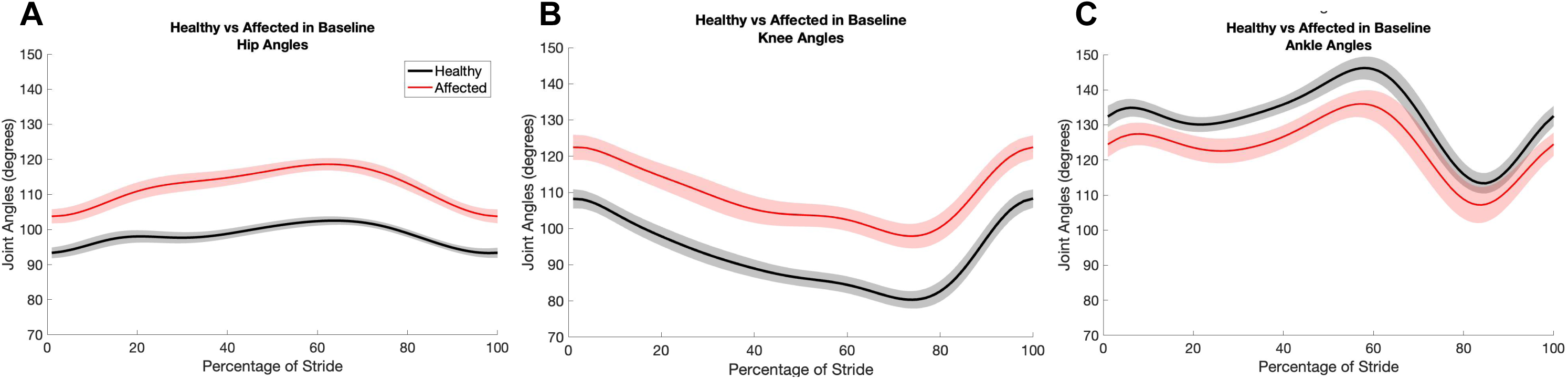
Joint angle range of motion (ROM) comparison for Healthy and Affected pigs. A) ROM for hip, B) knee, and C) ankle joints during walking test. (n= 2-5/group).

**Table 1.** Gait Analysis for Healthy and Affected pigs.

|  | Healthy | Affected |  | Healthy | Affected |
| --- | --- | --- | --- | --- | --- |
| *Hip ROM (degrees)* | 10.11 | 16.85 | *SD Swing Time (seconds)* | 0.019 | 0.021 |
| *Hip ROM SD (degrees)* | 0.821 | 1.642 | *Mean Stance Time (seconds)* | 0.449 | 0.371 |
| *Knee ROM (degrees)* | 29.54 | 27.03 | *SD Stance Time (seconds)* | 0.039 | 0.041 |
| *Knee ROM SD (degrees)* | 2.170 | 2.806 | *Mean Step Time (seconds)* | 0.745 | 0.631 |
| *Ankle ROM (degrees)* | 33.90 | 30.19 | *SD Step Time (seconds)* | 0.040 | 0.044 |
| *Ankle ROM SD (degrees)* | 3.178 | 4.960 | *Mean Stance Percent (percent)* | 0.601 | 0.586 |
| *Mean Swing Time (seconds)* | 0.296 | 0.260 | *SD Step Time (seconds)* | 0.028 | 0.036 |

Gait analysis data for Healthy and Affected groups at 8-9 months of age. Values are means, n=2-5/group.

### In Vivo Muscle Function

At 9 months of age (n=2-5), flexors in Affected pigs had reduced absolute maximum torque compared to Healthy pigs (**Figure 4A**, 48%, 95% CI -1.86 to -0.61). Fatigue curves for Affected appeared more linear and reduced compared to Healthy (**Figure 4B**). Area under the curve for flexor torque was 58% lower (95% CI -55.56 to -20.51) (**Figure 4C**). In addition, the fatigue index was 33% greater (95% CI -36.25 to 96.71) in Affected vs. Healthy (**Figure 4D**). For extensor muscles, maximum torque and fatigue metrics were similar between groups (**Figure 4 E-H**).

**Figure 4.**
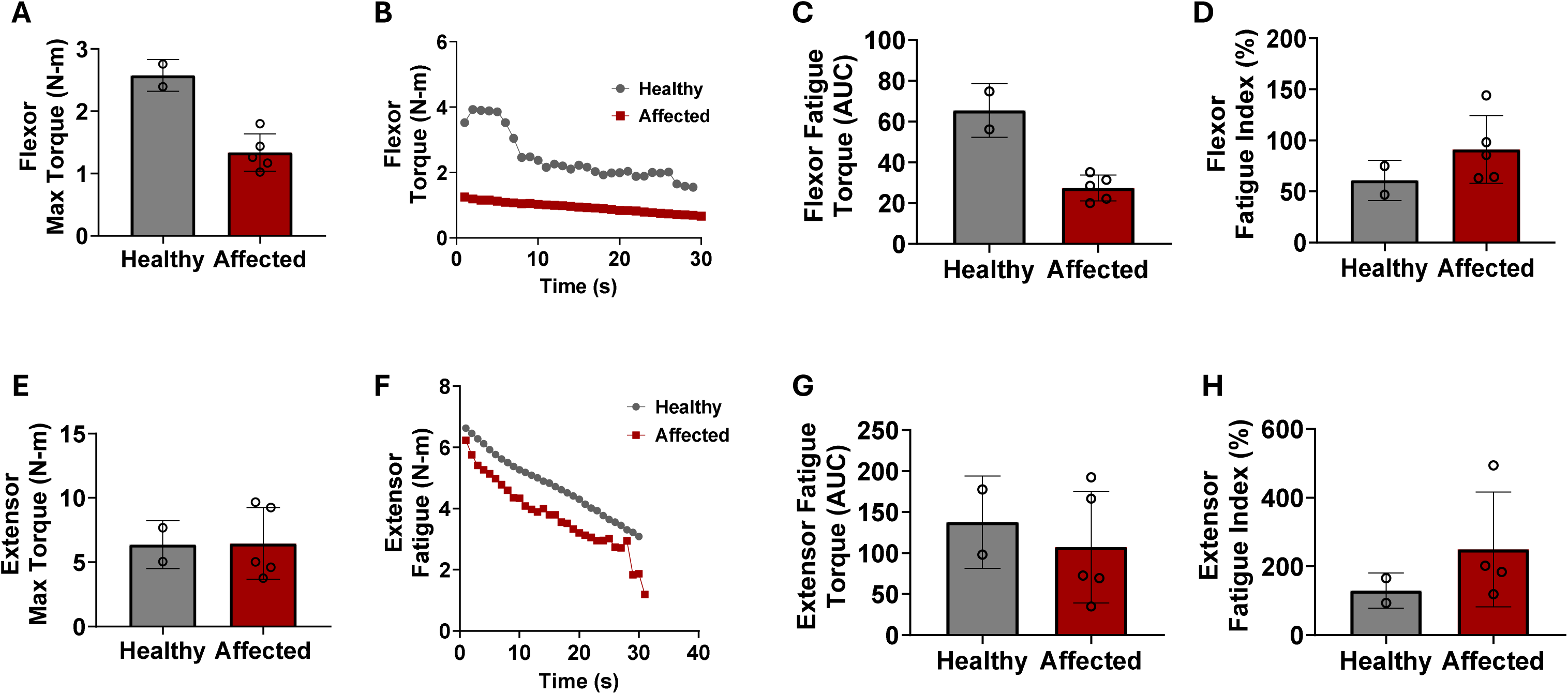
Muscle torque from Healthy and Affected flexor and extensor muscles. A) peak flexor torque, B) flexor fatigue curves, C) flexor fatigue Area Under the Curve (AUC), D) flexor fatigue index. E) peak extensor torque, F) extensor fatigue curves, G) extensor fatigue Area Under the Curve (AUC), H) extensor fatigue index. (n= 2-5/group).

### Muscle Fiber Histopathology

Muscle fiber CSA was determined in cranial tibial (flexor) and gastrocnemius (extensor) muscles (**Figure 5**). CSA was partitioned into 2000 µm^2^ increments, creating relative frequency histogram plots for Healthy and Affected pigs (**Figure 5A**). Flexor muscles of BMD pigs shifted the CSA frequency curve to the left (**Figure 5A, B**). There were 43% (95% CI 17.74 to 57.38) more small fibers (≤6000 µm^2^) and a 75% (95% CI -55.14 to -15.66) reduction in large fibers (≥8000 µm^2^) (**Figure 5C, D**). However, extensor muscle fiber CSAs appeared similar across fiber sizes (**Figure 5E-H**).

**Figure 5.**
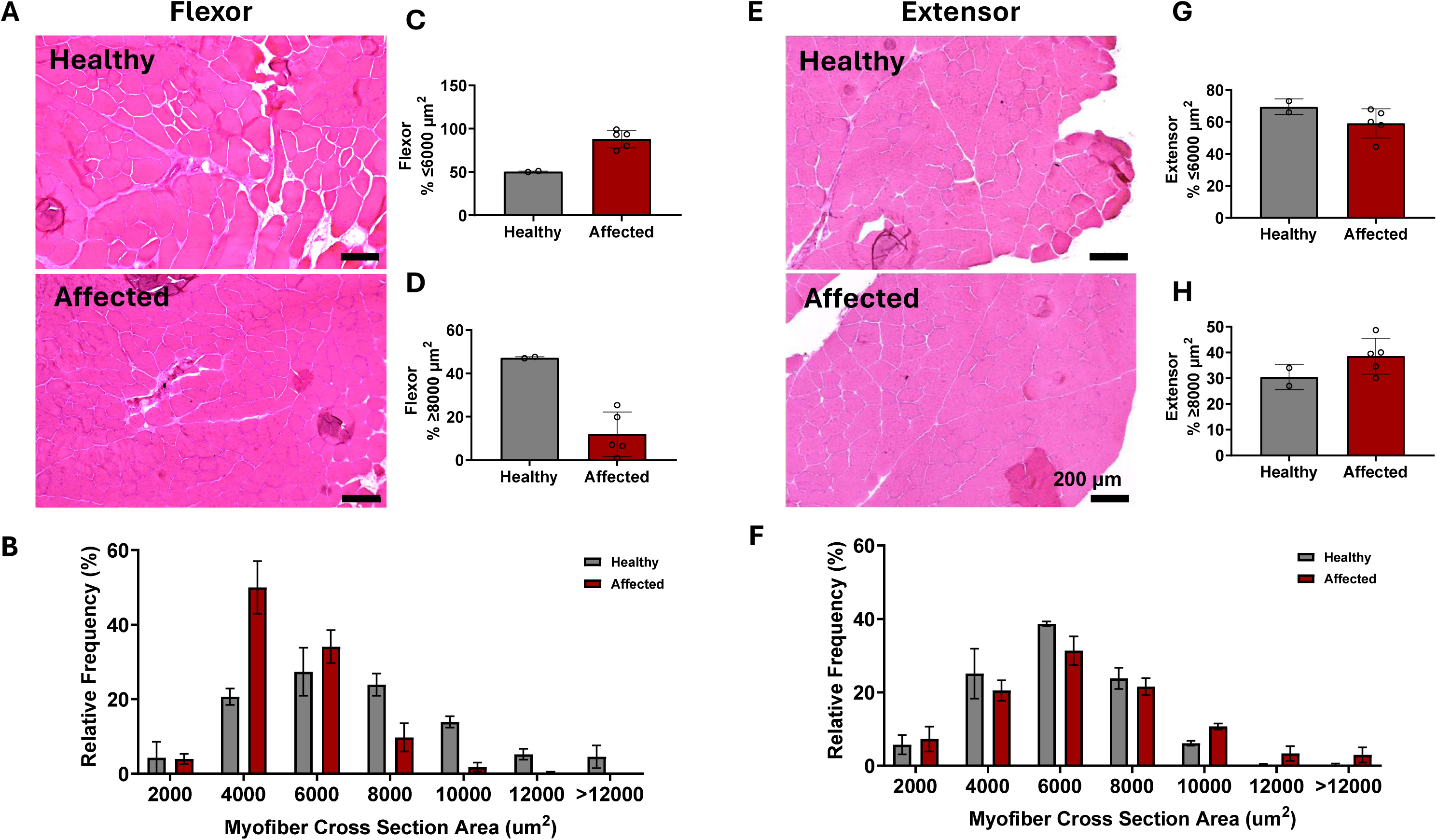
Histology of flexor and extensor muscles of Healthy and Affected pigs. A) Representative images of Healthy and Affected extensor muscle cross sections. Scale bars = 200 µm. B) Frequency histogram from binned cross-sectional areas (CSA) of extensor muscles. C) Summary data of small and D) large diameter fibers from extensor muscles. E) Representative images of Healthy and Affected flexor muscle cross sections. Scale bars = 200 µm. F) Frequency histogram from binned (0 - >12,000 in 2,000 µm^2^ increments) cross-sectional areas (CSA) of flexor muscles. G) Summary data of small and H) large diameter fibers from flexor muscles. (n= 2-5/group)

### Dystrophin Expression

Relative dystrophin protein decreased by 34% (95% CI -87.9 to 20.2) in cranial tibial muscles (flexors) and 37%, with significant variability, (95% CI -138.1 to 62.73) in gastrocnemius (extensors) of Affected pigs versus Healthy (**Figure 6A, B**, n=2-5/group).

**Figure 6.**
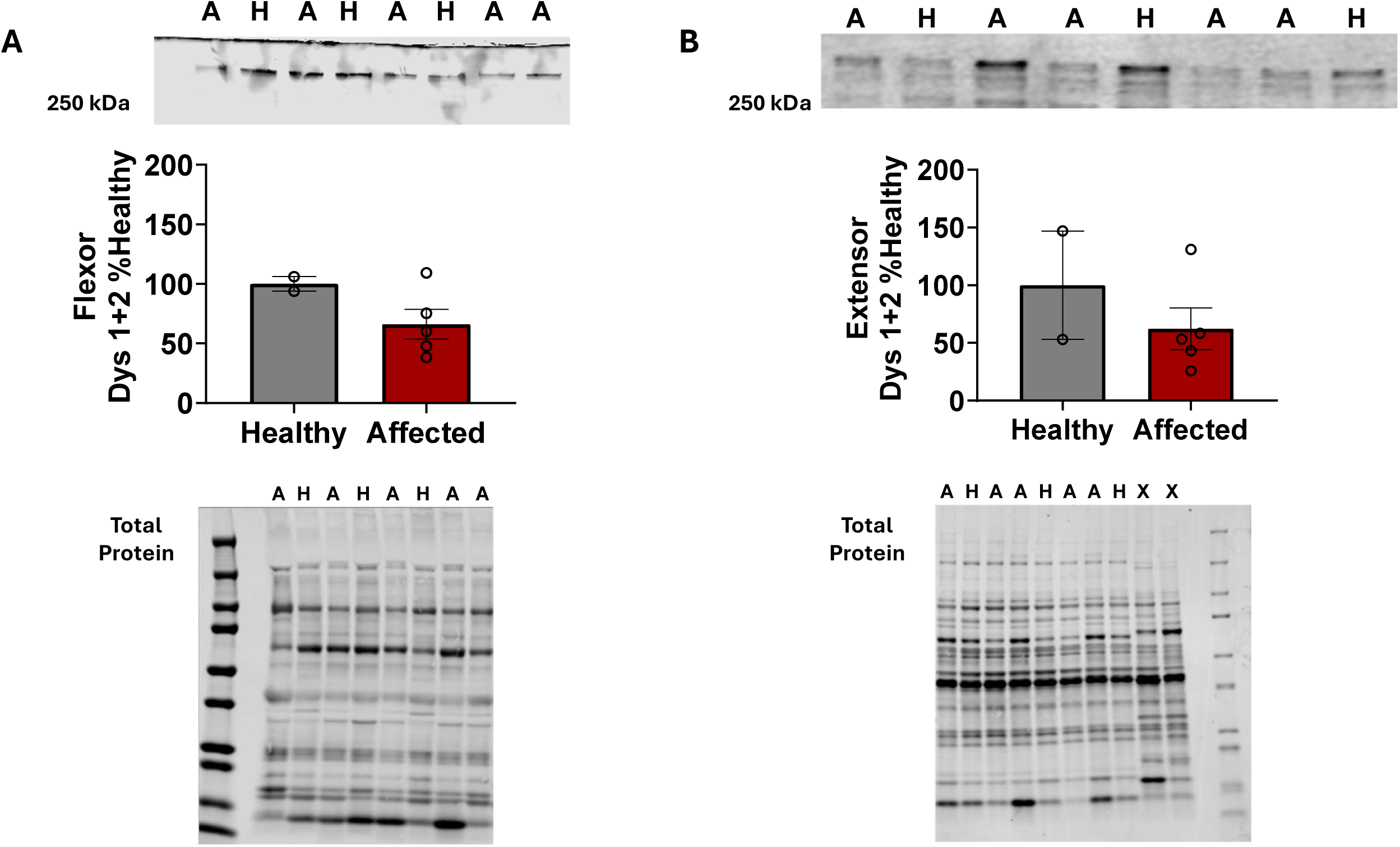
Dystrophin 1 and 2 expression in Affected compared to Healthy pigs reported as percent of Healthy. A) Representative blot image at 427 kDa (top), mean densiometric data (middle), and total protein (bottom) for A) Extensor muscles and B) Flexor muscle. (n= 2-5/group).

### Correlation of Dystrophin Expression to Functional Measures

Linear regression was used to discern whether there was a correlation between relative dystrophin protein abundance from flexor and extensor muscles of Healthy and Affected pigs to markers of muscle function (**Figure 7, Supplementary Figure 1-3**) (n=2-5). Flexor torque was significantly correlated with dystrophin expression (**Figure 7A**. P=.049) while fatigue index was inversely correlated with relative dystrophin abundance in flexor muscles (**Figure 7B**. P =.013) but not extensor muscles (**Figure 7D-F**).

**Figure 7.**
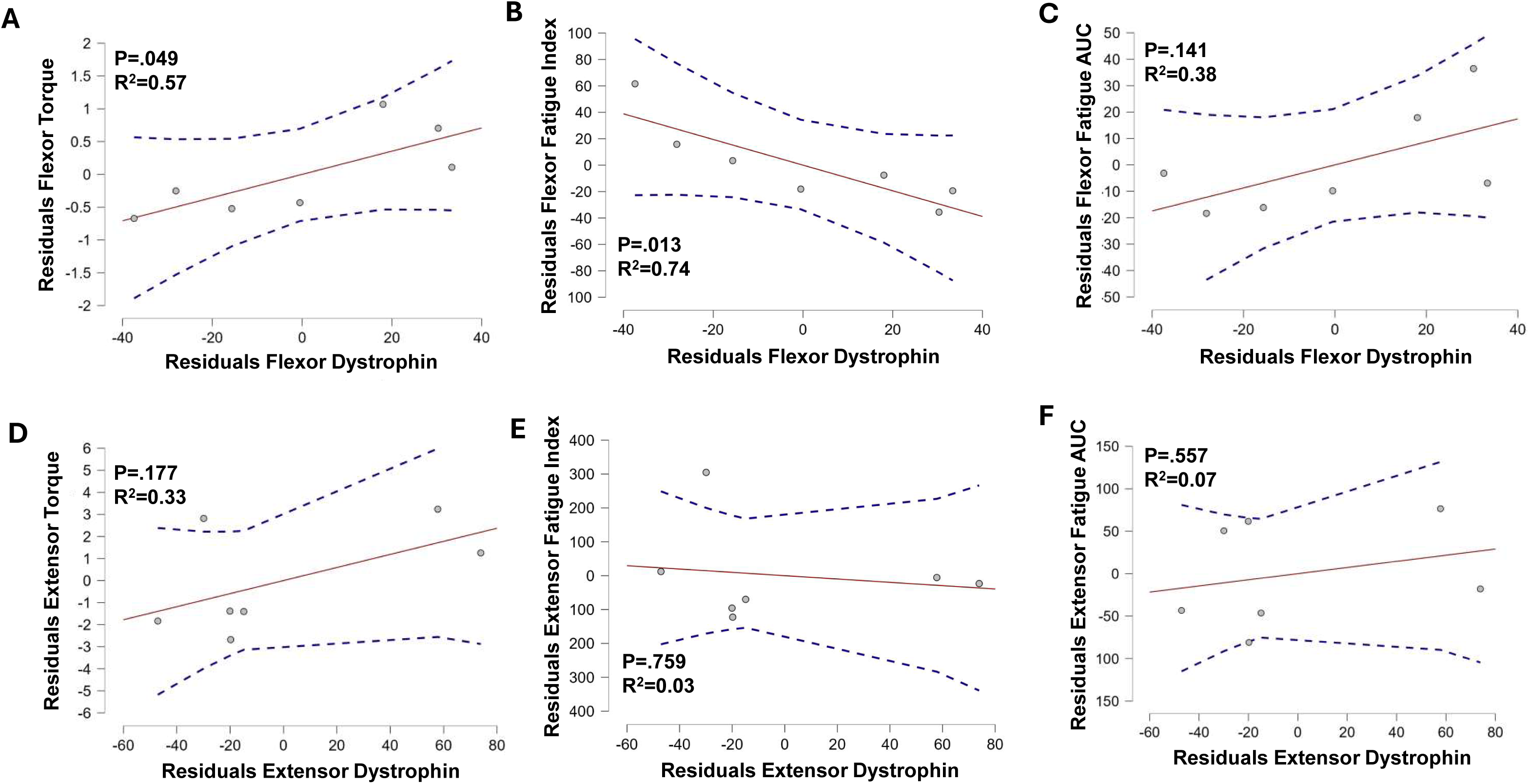
Correlation of Dystrophin protein expression to direct muscle function from Healthy and Affected pigs. *Flexor Muscles* A) Torque and B) fatigue index are significantly correlated with Dystrophin expression while the C) areas under fatigue curves (AUC) were not. *Extensor Muscles* D-F) None of the functional measures in extensor muscles correlated with dystrophin expression.

### Blood Panel Analysis

A comprehensive blood chemistry panel revealed that CK concentration was increased by 250%, (95% CI -215 to 1973) in Affected pigs compared to Healthy pigs. All other measured circulating factors were similar between groups (**Table 2**, n=3-5).

**Table 2.** Blood panel for Healthy and Affected pigs.

| Blood Marker | Healthy | Affected | Blood Marker | Healthy | Affected |
| --- | --- | --- | --- | --- | --- |
| *Creatine Kinase (pg/dL)* | 558.3 | 1438.0 | *ALKP (U/L)* | 70.0 | 76.3 |
| *Total Protein (g/dL)* | 7.4 | 7.0 | *AST (U/L)* | 70.0 | 109.4 |
| *Albumin (g/dL)* | 4.8 | 4.3 | *ALT (U/L)* | 46.5 | 46.4 |
| *Calcium (mg/dL)* | 12.5 | 11.6 | *Globulins (g/dL)* | 2.7 | 2.7 |
| *Phosphorous (mg/dL)* | 9.0 | 8.8 | *Albumin: Globulin Ratio* | 1.8 | 1.6 |
| *Glucose (mg/dL)* | 138.5 | 102.6 | *Sodium (mEq/L)* | 180.5 | 160.6 |
| *BUN (mg/dL)* | 21.0 | 18.6 | *Potassium (mEq/L)* | 6.1 | 10.4 |
| *Creatinine (mg/dL)* | 1.5 | 1.4 | *Chloride (mEq/L)* | 133.0 | 117.2 |
| *Bilirubin - total (mg/dL)* | 0.1 | 0.1 | *Sodium: Potassium Ratio* | 29.7 | 20.7 |

Blood panel data for Healthy and Affected groups at 8-9 months of age. Values are given in picograms (pg), milligrams (mg), or grams (g) per deciliter (dL), or units (U) or milliequivalents (mEq) per liter (L). Values are means, n=2-5/group.

## Discussion

In this pilot study, we present a phenotypic characterization of a naturally occurring porcine model of BMD, focusing specifically on hindlimb skeletal muscles and gait analysis. This porcine model may fill a critical gap in translational research as large animal models of BMD are limited. These findings build upon our group’s previous work, investigating the functional, histological, and biochemical markers to determine the validity in recapitulating the phenotype of human BMD patients (Hollinger et al., 2014; Selsby et al., 2015). Moreover, given the increasing focus on gene and exon skipping therapies aimed at converting DMD into a milder BMD phenotype (Nakamura, 2019), the population of BMD and BMD-like presentations in humans may rise.

Among the most significant findings of our pilot study were the potential gait abnormalities seen in Affected pigs, specifically hip ROM. Standard deviation from all gait measures demonstrate an almost 30% greater inconsistency in Affected pigs, while overall spatiotemporal parameters and joint angles of the gait cycle showed modest deviations, the 40% (95% CI -0.64 to 14.12) increase in hip ROM mirrors key biomechanical phenotypes observed in human BMD patients where compensatory hip movements are required to maintain forward propulsion due to weakness in the proximal leg muscles (Gao et al., 2025). These findings suggest that the hindlimb musculature in Affected pigs may exhibit functional maladaptation to underlying muscle weakness like human patients (Hou, Du, & Wu, 2022; Stirm et al., 2021). The lack of observed differences in knee and ankle ROM may reflect early or milder disease progression, typically seen in BMD where distal muscles are initially spared (Thada, Bhandari, Forshaw, & Umapathi, 2025). These gait alterations further suggest this model may recapitulate the human BMD phenotype (Nakamura et al., 2023) reinforcing the translational potential in evaluating functional endpoints for therapeutic intervention. Importantly, these kinematic disruptions provide a non-invasive, longitudinal marker that can be leveraged to monitor disease progression and response to treatment.

Muscle function testing and histopathological analyses further suggested the presence of a Becker-like phenotype in Affected pigs. *In vivo* muscle torque assessments revealed reductions in the flexor muscles in Affected pigs compared to Healthy, with no evident changes in extensor function. This asymmetry may suggest differential muscle involvement based on anatomical usage of a quadruped and disease burden, aligning with prior studies indicating a heterogenous pathology across muscle groups in BMD (Bello et al., 2016; Comi et al., 1994). Histopathological analysis of muscle CSA aligned with cycles of degeneration and regeneration common to dystrophinopathies (Darras, Urion, & Ghosh, 2022), a spectrum of X-linked muscle diseases. While the extensor muscles of Healthy and Affected showed no difference in muscle fiber morphology, the flexor muscles of Affected pigs displayed a leftward shift in fiber size distribution, with increases in the percentage of small muscle fibers and reductions in the percentage of large muscle fibers. This change in muscle fiber size is consistent with muscle injury and repair frequently observed in dystrophinopathies indicating ongoing cycles of degenerative and regenerative activity (Ripolone et al., 2022). Furthermore, average CK levels in blood were 2.5-fold greater in Affected pigs, a common biomarker of muscle fiber membrane instability and damage. Taken together, these findings suggest the BMD pig model exhibits functional deficits and histopathological hallmarks of disease in select muscles, similar to observations in humans with BMD.

Analysis of dystrophin revealed moderate reductions in both extensor and flexor hindlimb muscles of Affected pigs. Although this reduction in dystrophin is less severe than previous investigations into the diaphragm, longissimus, and psoas (Selsby et al., 2015), these values are consistent with the in-frame nature of the mutation in Exon 41, likely permitting some dystrophin expression, but, given previous reports and data presented herein, not at levels sufficient to prevent disease pathology. The presence of dystrophin at reduced levels aligns with the intermediate clinical progression of BMD, where dystrophin is partially functional but unable to maintain normal muscle structure and function (Anthony et al., 2011; Bushby et al., 1993; Hoffman et al., 1989). Given the spectrum of severity noted in humans with BMD, modest reductions in dystrophin, when viewed alongside preliminary functional and histological deficits support the conclusion that this model may provide a translational perspective for therapeutic development. In addition, dystrophin expression was highly correlated with peak torque and fatigue index from flexor muscles. This may be due, at least in part, to a greater reliance on intermediate/slower, more fatigue resistant muscle fiber types and a reduction in the contribution of fast fatigable fibers to functional measures in peak torque concomitant with declines in dystrophin as demonstrated by others (Webster et al., 1988).

Limitations to the current study were that we were only able to study eight pigs (5 with BMD and 3 healthy from the same background). Although the power of such a study is low, observations in Affected pigs were comparable to recent published results in human BMD patients.

## Conclusion

This pilot study provides a gait characterization of a naturally occurring porcine model of BMD with an in-frame Exon 41 mutation resulting in dystrophin insufficiency. Our findings indicate a Becker-like phenotype including disease mediated alterations in hip ROM, reduced flexor muscle function and muscle fiber CSA, and elevated CK concentrations. Dystrophin expression was inversely correlated with fatigue index in flexor muscles. In contrast, extensor muscles did not display a Becker-like phenotype across all parameters, suggesting heterogeneity in muscle pathology. These abnormalities closely mirror the human BMD phenotype, particularly in functional and histopathological features of the hindlimb musculature. Together, these results further suggest the BMD pig as a translationally relevant large animal model for studying disease progression and evaluating the safety and efficacy of emerging therapeutic advancements.

## Data Availability Statement

The data that supports the findings of this study are available at the following link. https://doi.org/10.7910/DVN/ZOX424.

## Author Contributions

Conceptualization: A.B.M., P.P.N., Data Curation: A.B.M., J.A.K., S.G., J.M.Y., P.P.N., Formal Analysis: A.B.M., J.A.K., S.G., J.M.Y., P.P.N., Funding Acquisition: A.B.M., P.P.N., Investigation: A.B.M., J.A.K., M.M., S.G., J.T.S., P.P.N., Methodology: A.B.M., M.M., K.C., J.M.Y., P.P.N., J.T.S., J.W.R., Z.E.K., Project Administration: A.B.M., P.P.N., Resources: A.B.M., Z.E.K., J.W.R., J.M.Y., J.T.S., N.E.P.D., P.P.N., Validation: A.B.M., P.P.N., Visualization: A.B.M., P.P.N., Writing – original draft: A.B.M., J.A.K., Writing – review and editing: A.B.M., J.A.K., J.M.Y., P.P.N., J.T.S., All authors have read and approved the final version of this manuscript and agree to be accountable for all aspects of the work ensuring that questions related to accuracy or integrity of any part of the work are appropriately investigated and resolved. All persons designated as authors qualify for authorship and all those who qualify for authorship are listed.

## Acknowledgements

We would like to thank Veterinary Medical Park at Texas A&M University (TAMU) for their husbandry and veterinary care. We would also like to thank Eden Haneline for her assistance in tracking videos for analysis.

## Figure Legends

**Supplementary Figure 1.**
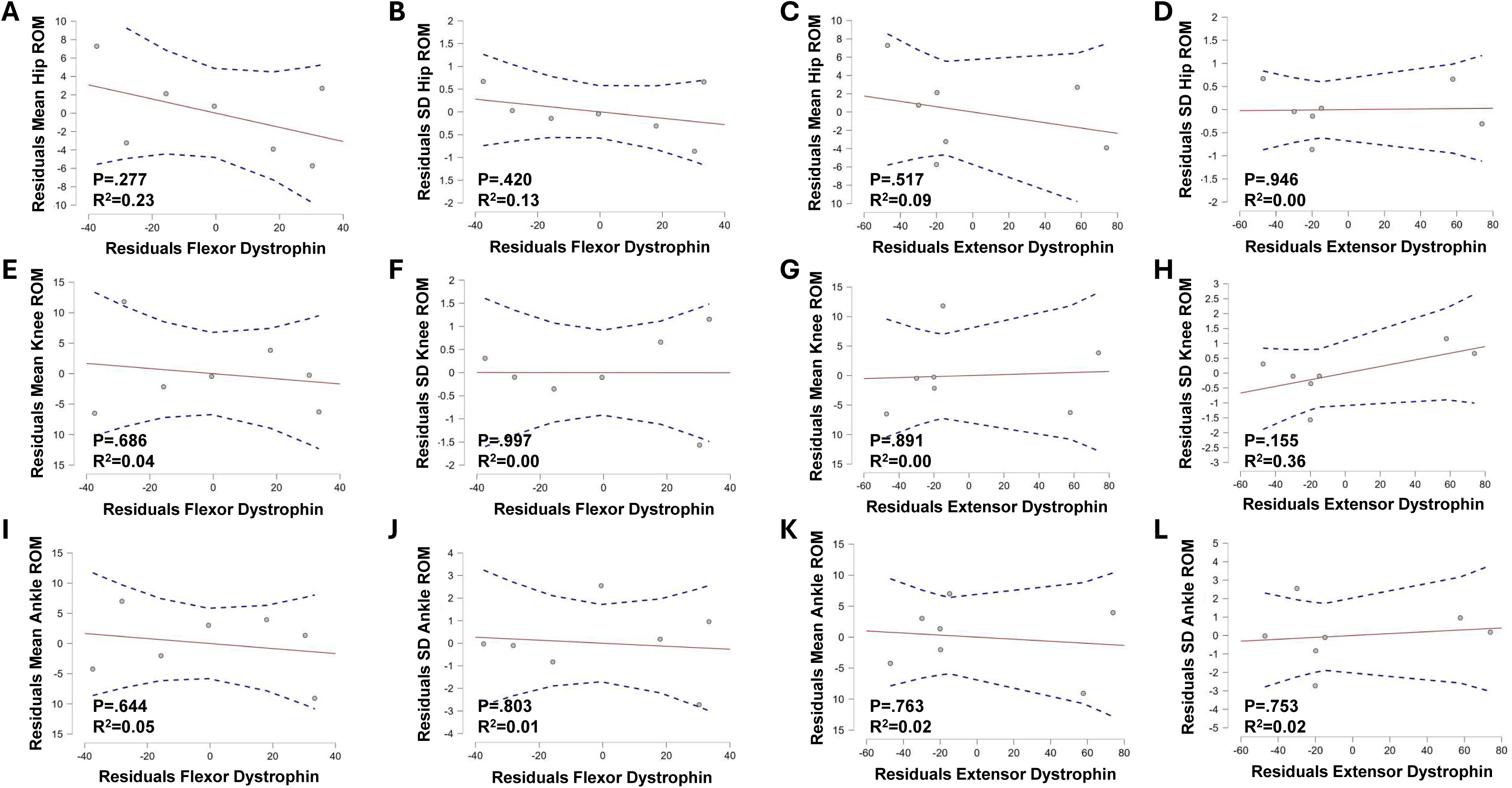
Correlations of Dystrophin protein expression from Healthy and Affected extensor and flexor muscles to joint range of motion (ROM) variables. *Flexor Dystrophin* A, B, E, F, I, J) joint ROM variables were not well correlated with dystrophin expression. In addition, dystrophin expression in *Extensor Muscles* C, D, G, H, K, L) was also not well correlated with ROMs.

**Supplementary Figure 2.**
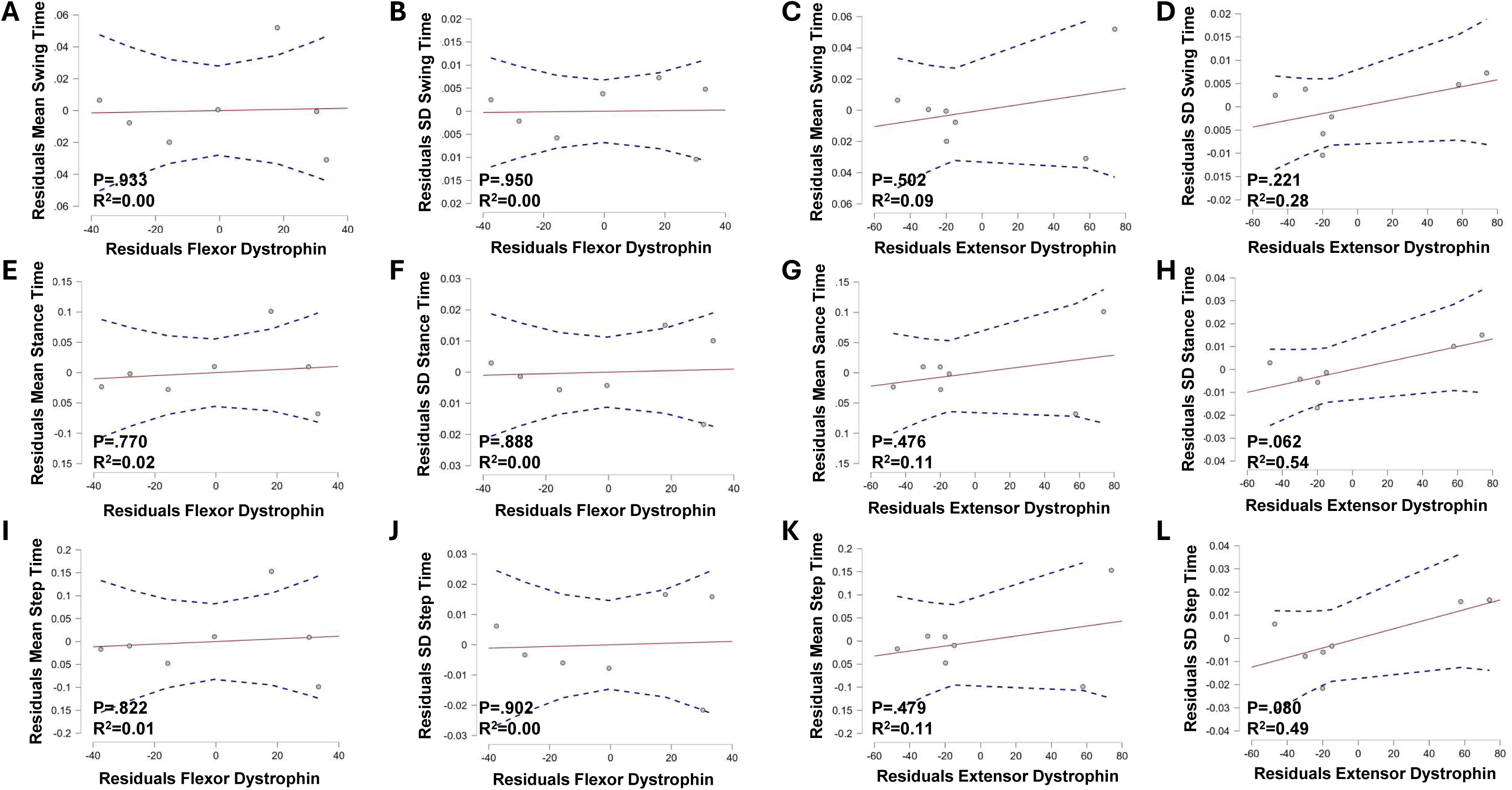
Correlations of Dystrophin protein expression from Healthy and Affected extensor and flexor muscles to temporal variables in the gait cycle. *Flexor Dystrophin* A, B, E, F, I, J) temporal variables were not well correlated with dystrophin expression. In addition, *Extensor Muscles* C, D, G, H, K, L) were also not well correlated.

**Supplementary Figure 3.**
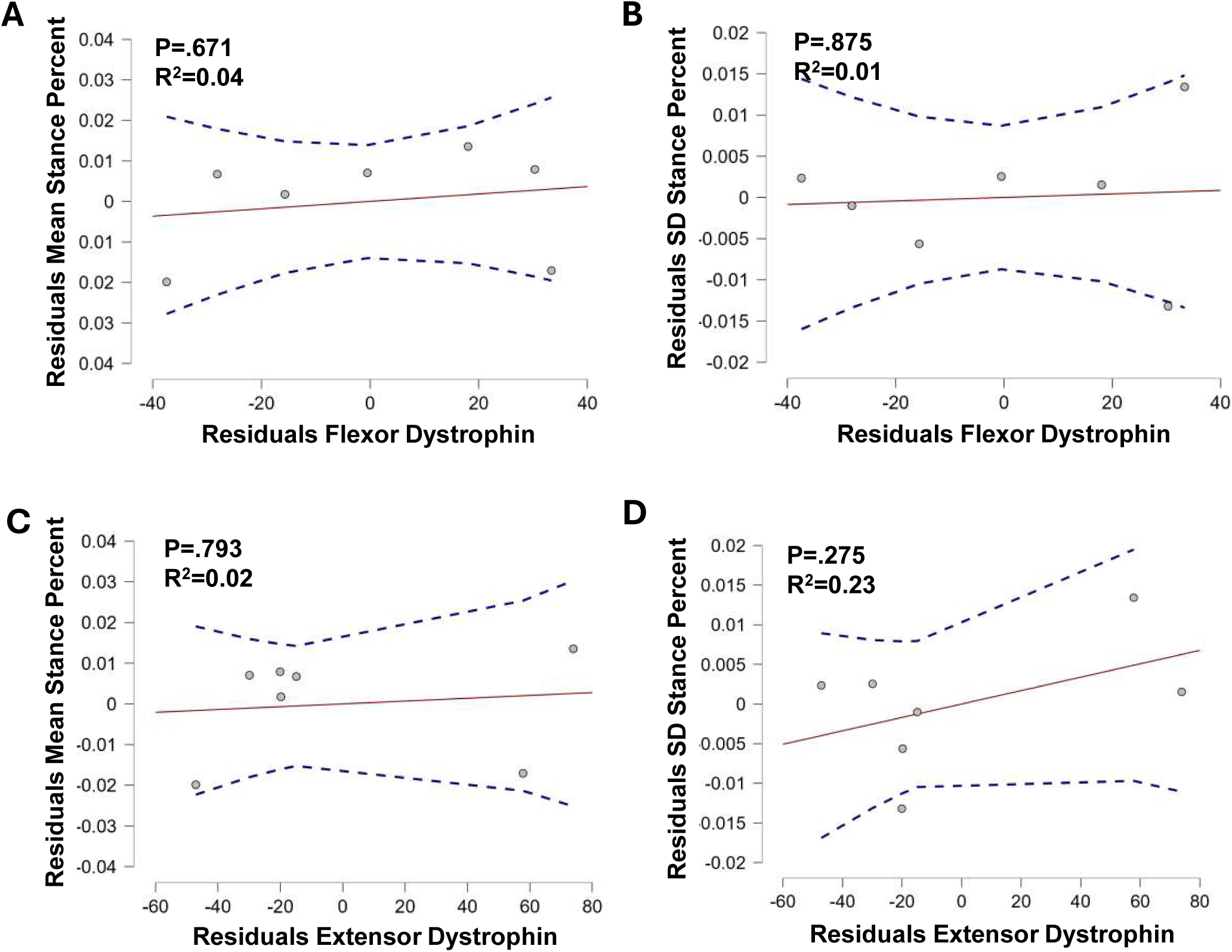
Evaluation of stance percent parameters compared to dystrophin expression in either flexor (A and B) or extensor (C and D) muscles.

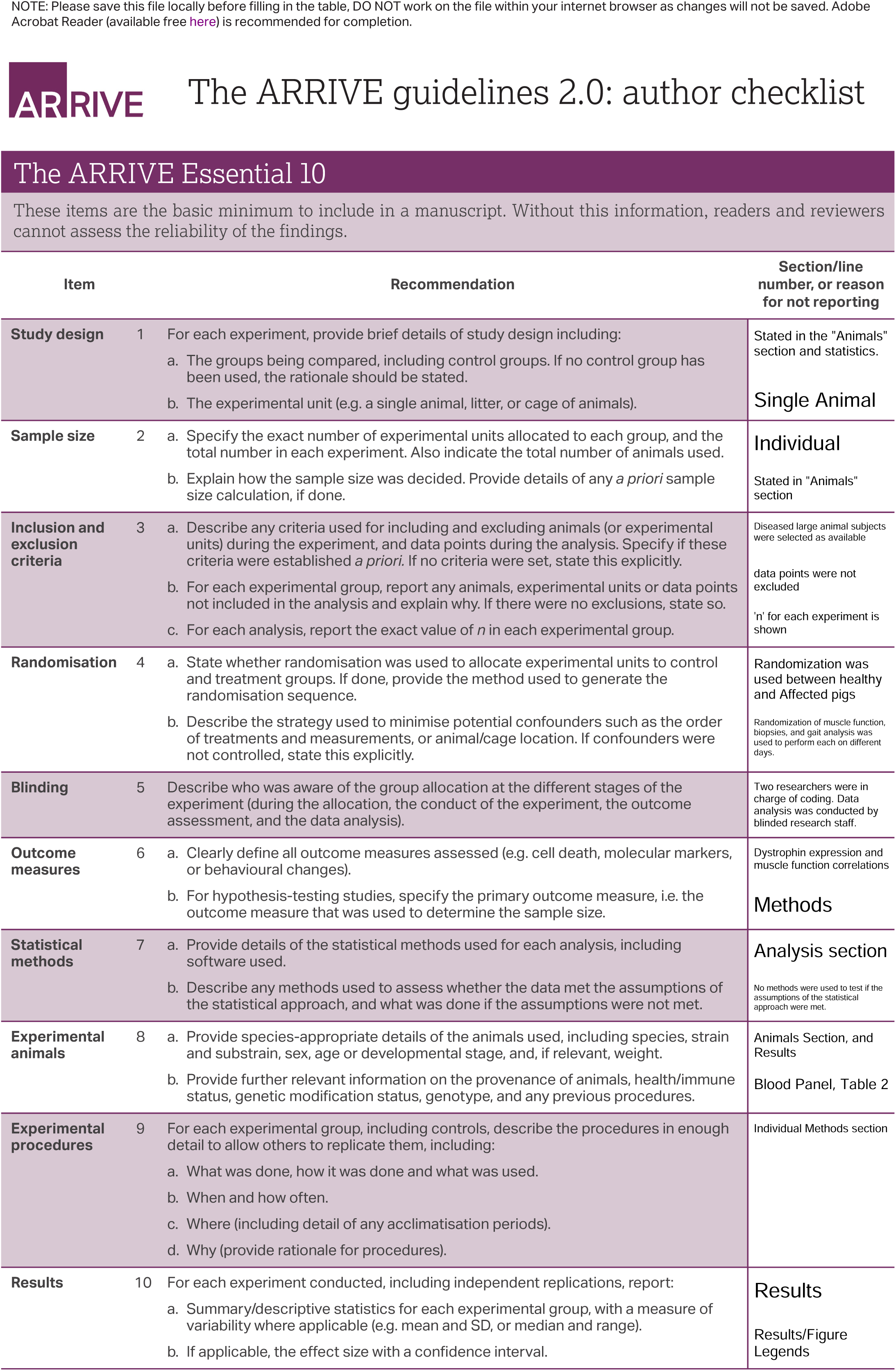

